# Primate immunodeficiency virus Vpx and Vpr counteract transcriptional repression of proviruses by the HUSH complex

**DOI:** 10.1101/293001

**Authors:** Leonid Yurkovetskiy, Mehmet Hakan Guney, Kyusik Kim, Shih Lin Goh, Sean McCauley, Ann Dauphin, William Diehl, Jeremy Luban

## Abstract

Drugs that inhibit HIV-1 replication and prevent progression to AIDS do not eliminate HIV-1 proviruses from the chromosomes of long-lived CD4^+^ memory T cells. To escape eradication by these antiviral drugs, or by the host immune system, HIV-1 exploits poorly defined host factors that silence provirus transcription. These same factors, though, must be overcome by all retroviruses, including HIV-1 and other primate immunodeficiency viruses, in order to activate provirus transcription and produce new virus. Here we show that Vpx and Vpr, proteins from a wide range of primate immunodeficiency viruses, activate provirus transcription in human CD4^+^ T cells. Provirus activation required the DCAF1 adaptor that links Vpx and Vpr to the CUL4A/B ubiquitin ligase complex, but did not require degradation of SAMHD1, a well-characterized target of Vpx and Vpr. A loss-of-function screen for transcription silencing factors that mimic the effect of Vpx on provirus silencing identified all components of the Human Silencing Hub (HUSH) complex, FAM208A (TASOR/RAP140), MPHOSPH8 (MPP8), PPHLN1 (PERIPHILIN), and MORC2. Vpx associated with the HUSH complex components and decreased steady-state levels of these proteins in a DCAF-dependent manner. Finally, *vpx* and FAM208A knockdown accelerated HIV-1 and SIV_MAC_ replication kinetics in CD4^+^ T cells to a similar extent, and HIV-2 replication required either *vpx* or FAM208A disruption. These results demonstrate that the HUSH complex restricts transcription of primate immunodeficiency viruses and thereby contributes to provirus latency. To counteract this restriction and activate provirus expression, primate immunodeficiency viruses encode Vpx and Vpr proteins that degrade HUSH complex components.

---

When provided *in trans*, many primate immunodeficiency virus Vpx and Vpr orthologues increase HIV-1 reverse transcription and transduction efficiency in dendritic cells, macrophages, and resting CD4^+^ T cells (Baldauf et al., 2012; Goujon et al., 2006; Lim et al., 2012; Sharova et al., 2008; Srivastava et al., 2008). As substrate adaptor proteins for the DCAF1-CUL4A/B E3 ubiquitin ligase, Vpx and Vpr increase the concentration of deoxynucleotide triphosphate (dNTP) levels in target cells by degrading the deoxynucleotidetriphosphate (dNTP) hydrolase SAMHD1 (Hrecka et al., 2011; Laguette et al., 2011; Lim et al., 2012). Nonetheless, Vpx and Vpr have additional effects on expression of transduced reporter genes that are not explained by SAMHD1 degradation or by increase in dNTP concentration (Goh et al., 1998; Miller et al., 2017; Pertel et al., 2011a; Reinhard et al., 2014).

To better understand the effect on provirus reporter gene expression, *vpx* was introduced before, during, or after transduction of a reporter gene (Fig. 1a). Jurkat CD4^+^ T cells were transduced with a dual-promoter, lentiviral vector that expresses codon-optimized SIV_MAC_251 *vpx* from the spleen focus forming virus (SFFV) promoter and puromycin acetyltransferase (puro^R^) from the PPIA (CypA) promoter (Neagu et al., 2009; Reinhard et al., 2014) (Lenti 1 in the Fig. 1a time-line, Supplementary Fig. 1a, and Supplementary Table 1). A control Lenti 1 vector was used that lacks *vpx* (Supplementary Fig. 1a). Puromycin was added to the culture on day three to select those cells that had been transduced with Lenti 1. On day seven, cells were transduced with a second lentivector bearing a codon-optimized *gag*-*gfp* reporter gene expressed from the SFFV promoter, as well as SIV_MAC_251 *vpx* expressed from the CypA promoter (Lenti 2 in the Fig. 1a timeline and Supplementary Fig.1a). A control Lenti 2 vector was used that lacks *vpx* (Supplementary Fig.1a). On day ten, virus-like particles (VLPs) containing Vpx protein were added to the twice-transduced cells (Fig. 1a). As controls, VLPs lacking Vpx were used, or no VLPs were added. On day fourteen, the percent GFP^+^ cells under each condition was assessed by flow cytometry using standard gating for viable, singlet, lymphoid cells (Supplementary Figure 1b). Vpx increased the percentage of GFP^+^ cells, whether *vpx* was transduced before, or concurrent with, reporter gene transduction, or if Vpx protein was delivered by VLPs after reporter gene transduction (Fig. 1b and Supplementary Fig 1c; n=3 biological replicates, p<0.02, 1-way ANOVA with Dunnett post-test). These results suggest that the transduced reporter gene was actively silenced and that *vpx* overcame reporter silencing.

**Figure 1.**
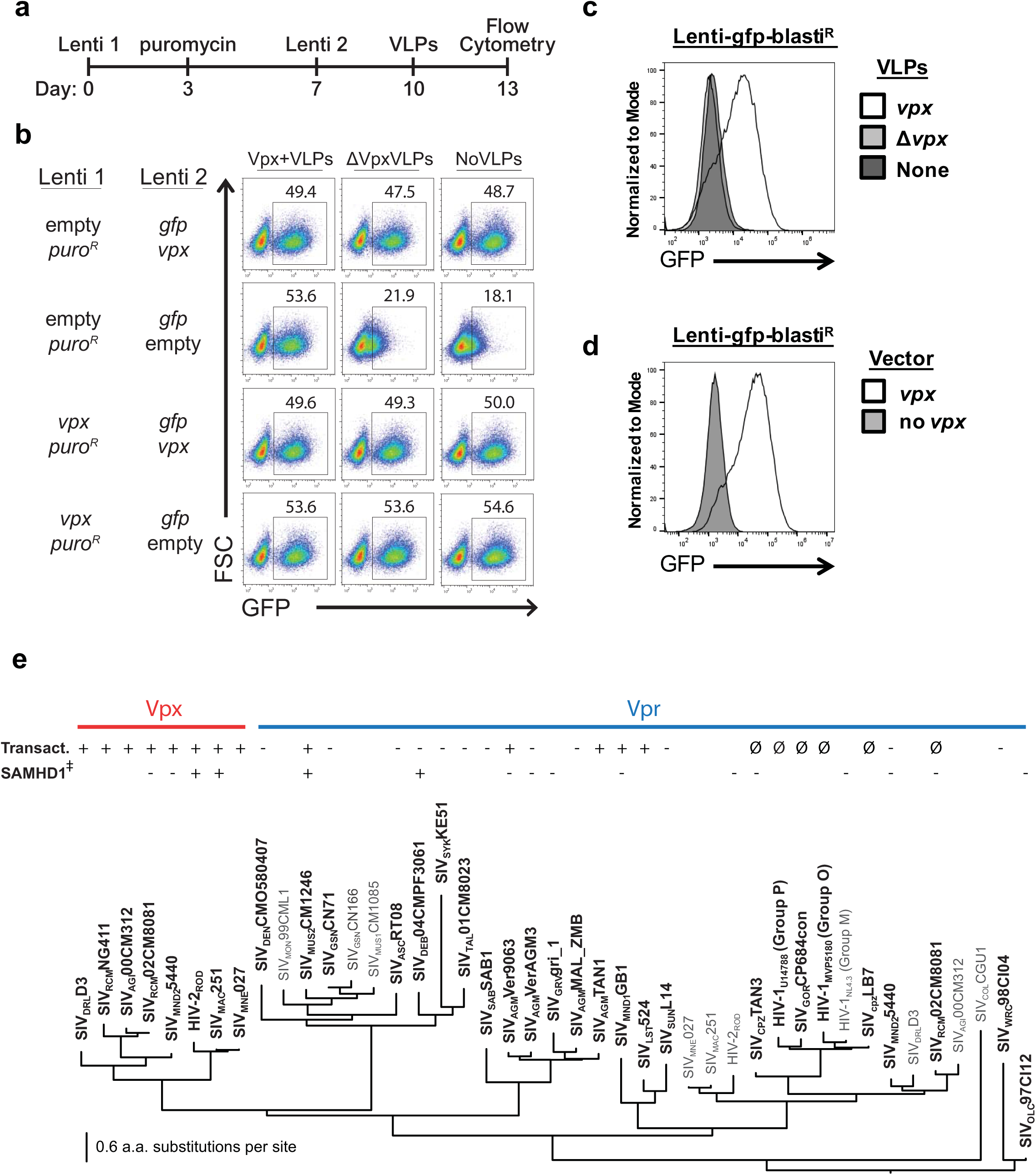
Diverse primate immunodeficiency virus *vpx* and *vpr* orthologues activate provirus transcription, whether delivered before, during, or after reporter provirus integration. **a,** Schematic of experimental protocol in (**b**). **b,** Flow cytometry plot showing percent GFP^+^ Jurkat cells after sequential transduction with the indicated lentivectors, followed by exposure to the indicated VLPs. **c,d,** Histogram of flow cytometry signal in Jurkat cells transduced with *gfp*-reporter virus, and either exposed to the indicated VLPs (**c**), or transduced with the indicated vectors (**d**). **e,** Phylogenetic tree showing evolutionary relationship of Vpx and Vpr proteins. The transactivation activity of Jurkat reporter lines, tested as in (**d**), and human SAMHD1 degradation activity (Lim et al., 2012), are indicated. ∅ indicates Vprs that were too toxic (G2 arrest) for assessment. All data shown is representative of at least three biological replicates.

To confirm that the findings in Fig. 1b were due to effects of *vpx* on transcriptional silencing of the reporter gene, and not due to effects on transduction efficiency, Jurkat T cells were first transduced with a vector in which the *gag*-*gfp* reporter gene was expressed from the SFFV promoter and blasticidin-S deaminase (blasti^R^) was expressed from the CypA promoter. Four days after transduction with the reporter vector and selection with blasticidin, cells were either challenged with Vpx^+^ VLPs, or transduced and selected with the dual-promoter lentivector encoding *vpx* and *puroR* (Lenti 1 in Supplementary Fig. 1a). Four days later the GFP signal was at background levels unless Vpx was provided, either by VLPs (Fig. 1c) or by *vpx* transduction (Fig. 1d). The effect of *vpx* on reporter gene expression was confirmed by qRT-PCR for the reporter mRNA (Supplementary Fig. 1d). Reporter gene silencing and reactivation by Vpx was not specific to the SFFV promoter since GFP signal was similar when the reporter gene was expressed from the human EEF1A1 (EF1α) promoter or from the Herpes simplex virus type 1 thymidine kinase (TK) promoter (Supplementary Fig. 1e). These results demonstrate that Vpx overcomes transcriptional silencing of the provirus.

To determine if the ability to activate transcription of silenced proviruses is peculiar to SIV_MAC_251 Vpx, representative Vpx and Vpr orthologues, selected from across the phylogeny of primate immunodeficiency viruses, were examined. All Vpx proteins tested, SIV_DRL_D3, SIV_RCM_NG411, SIV_AGI_00CM312, SIV_RCM_02CM8081, SIV_MND2_5440, HIV-2_ROD_, SIV_MAC_251, and SIV_MNE_027, had transactivating activity in human cells (Fig. 1e and Supplementary Fig 1f). Conservation of this activity in human cells among such divergent SIV orthologues was surprising given that SIV_RCM_NG411 Vpx and SIV_MND2_5440 Vpx do not degrade human SAMHD1, but they do degrade the SAMHD1 orthologue from their cognate primate host species (Lim et al., 2012). Several Vprs from SIVs that lack Vpx, including SIV_MUS2_CM1246, SIV_AGM_Ver9063, SIV_AGM_TAN1, SIV_MND1_GB1, and SIV_LST_524, also activated transcription of silent proviral reporters in human cells (Fig. 1e and Supplementary Fig. 1f). Results could not be obtained from this experimental system concerning the activity of Vprs encoded by SIV_CPZ_TAN3, HIV-1_U14788_ (Group P), SIV_GOR_CP684con, HIV-1_MVP5180_ (Group O), HIV-1_NL4-3_ (Group M), SIV_CPZ_LB7, and SIV_RCM_02CM8081, presumably because these orthologues caused cell cycle arrest and toxicity (Chang et al., 2004; Goh et al., 1998; He et al., 1995; Re et al., 1995) (indicated by ∅ in Fig. 1e). Vpx and Vpr sequence variability is among the highest observed for lentiviral coding sequences (McCarthy and Johnson, 2014); the sequences shown in Fig. 1e have an average amino acid identity of only 27%. Such diversity likely reflects rapidly evolving, host-pathogen interfaces (Fregoso et al., 2013), and precluded activity predictions based on amino acid sequence conservation to guide the engineering of loss-of-function mutations.

To gain insight into the mechanism by which Vpx overcomes transcriptional silencing of lentiviral transgenes, a loss-of-function screen was performed focusing on genes reported to contribute to silencing of retroviruses and other transcriptional targets (Chéné et al., 2007; Peterlin et al., 2017; Tchasovnikarova et al., 2015, 2017; Wang and Goff, 2017; Weinberg and Morris, 2016; Wolf and Goff, 2007). Jurkat T cells were transduced with lentivectors that confer puromycin resistance and express shRNAs (Pertel et al., 2011b) targeting either AGO1, AGO2, AGO3, DNMT3A, HDAC1, HP1, SUV39H1, SUV39H2, PIWIL2, TRIM28, SETDB1, FAM208A, MPHOSPH8, PPHLN1, or MORC2. After selection for five days with puromycin, cells were transduced with the Lenti 2 *gag-gfp* reporter vector without *vpx* (Supplementary Fig. 1a). Four days later, the change in expression of the *gfp* reporter due to the knockdowns was calculated as a percentage of the activity observed in a separate population of Jurkat cells transduced to express *vpx* (Fig. 2a). A given gene was implicated as a transcriptional silencing factor for the provirus reporter gene if the three shRNA targets for that gene differed significantly from that of the luciferase knockdown control (p<0.05, 1-way ANOVA with Dunnett post-test). shRNAs targeting each of the three core components of the Human Silencing Hub (HUSH) complex, FAM208A, MPHOSPH8, and PPHLN1, increased reporter gene expression (Fig. 2a).

**Figure 2.**
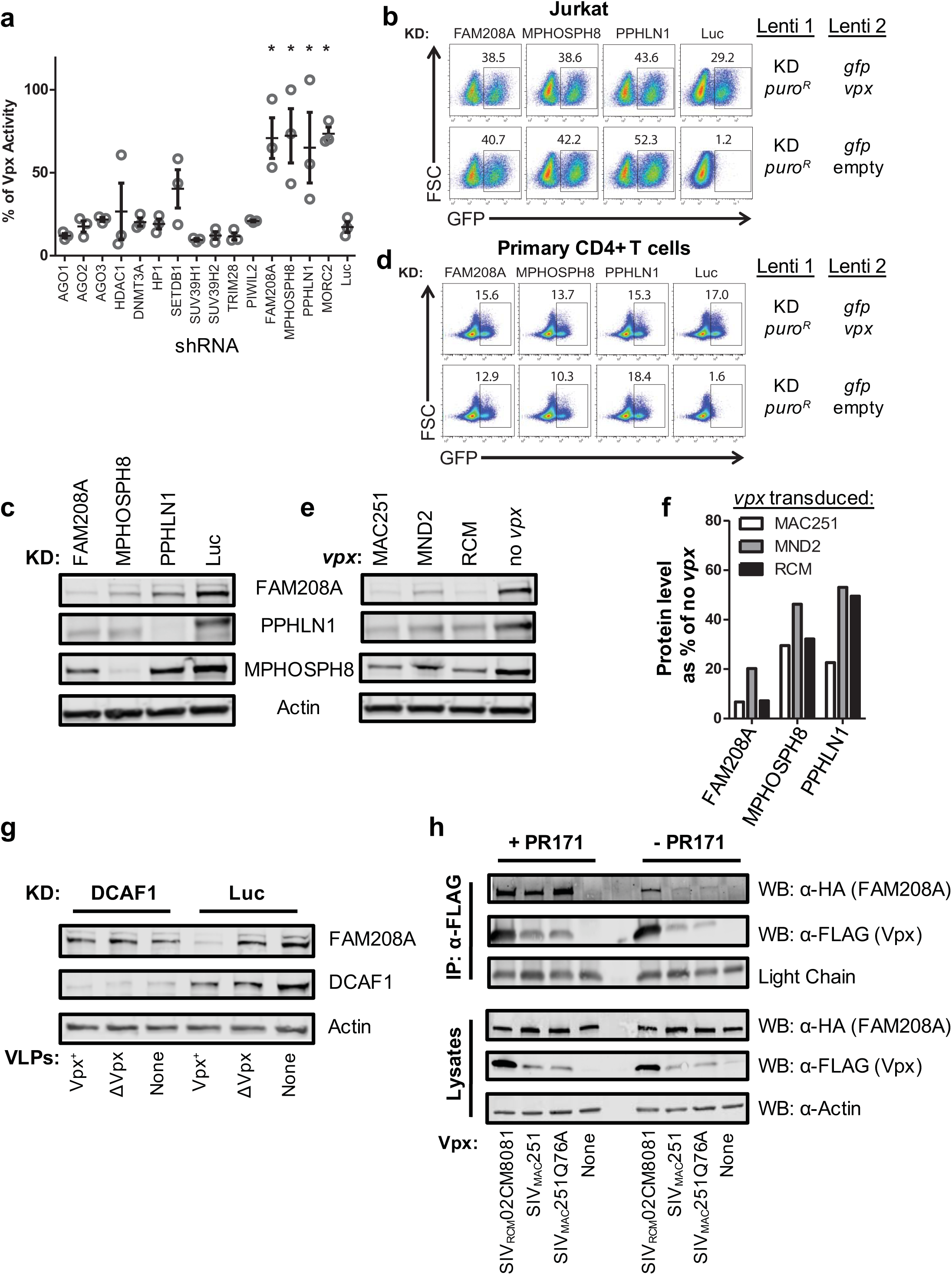
Vpx activates provirus transcription by degrading HUSH complex components. **a,** Jurkat cells transduced with shRNA-puro^R^ vectors targeting the indicated genes were selected with puromycin, transduced with Lenti 2-Δ*vpx*, and analyzed 5 days later. Plot depicts GFP signal in knockdown lines relative to Jurkats bearing SIV_MAC_251 *vpx* (mean ± S.E.M., n=3 shRNA target sites). *, *P*<0.05 as determined by 1-way ANOVA with Dunnett post-test, relative to luciferase knockdown control. **b,** Jurkat cells were transduced with the indicated shRNA-puro^R^ vectors and selected with puromycin. Resistant cells were transduced with *vpx*^+^ or Δ*vpx* Lenti 2 vector, and analyzed for GFP expression 7 days later. **c,** Immunoblot analysis for components of the HUSH complex in Jurkat cells expressing shRNA constructs used in (**b**). **d,** CD4^+^ T cells were activated for 3 days with PHA and then transduced and assayed as in (**b**). **e,** Immunoblot analysis of Jurkat lines transduced to express *vpx* from SIV_MAC_251, SIV_RCM_NG411, SIV_MND2_5440, or control. **f,** Levels of HUSH components in (**e**) shown as shRNA treated condition relative to control. **g,** FAM208A, DCAF1, and Actin immunoblot of Jurkat cells transduced with DCAF1 shRNA-puro^R^ vector or control, that were treated with Vpx^+^ or ΔVpx VLPs for 18 hrs. **h,** HEK293 cells were co-transfected with HA-FAM208A and the indicated FLAG-Vpx constructs. 18 hrs after transfection, cells were either exposed to proteasome inhibitor PR171 or left untreated. 8 hrs after inhibitor treatment cells were lysed, FLAG-Vpx was immunoprecipitated, and immunoblotted for FLAG-Vpx and HA-FAM208A. Immunoblotting of input lysates are shown below.

The effect on reporter gene expression in Jurkat T cells of the most effective shRNA target sequences for FAM208A, MPHOSPH8, and PPHLN1 is shown in Fig. 2b. The effectiveness of the knockdown of each of the HUSH complex components in Jurkat cells was confirmed by immunoblotting lysate from these cells with antibodies specific for FAM208A, PPHLN1, or MPHOSPH8 (Fig. 2c). As previously reported (Tchasovnikarova et al., 2015), knockdown of any individual HUSH complex component caused a decrease in the level of each of the other components. Similar results on reporter gene expression were obtained when FAM208A, MPHOSPH8, or PPHLN1 were knocked down in primary human CD4^+^ T cells (Fig. 2d). Knockdown of each of the HUSH complex components, then, had the same effect as *vpx* on lentiviral reporter gene expression (Fig. 2b and d and Supplementary Fig 2a). These results demonstrate that the HUSH complex is critical for provirus silencing and raise the possibility that Vpx acts as a substrate adaptor targeting HUSH components to DCAF1 and the CUL4A/B E3 ubiquitin ligase complex for degradation, in the same way that Vpx targets SAMHD1 (Hrecka et al., 2011; Laguette et al., 2011).

To determine if Vpx promotes the degradation of HUSH complex components, lysate from cells transduced to express SIV_MAC_251, SIV_MND2_5440, or SIV_RCM_NG411 *vpx* was immunoblotted with antibodies specific for FAM208A, PPHLN1, or MPHOSPH8. All three Vpx proteins reduced the steady-state level of all three core HUSH complex components (Fig. 2e). Among the three HUSH components, though, FAM208A protein levels were decreased more than the other two components (Fig. 2f) so ongoing experiments focused on the effect of Vpx on FAM208A. Indeed, in addition to the three Vpx proteins assessed in Fig. 2e, the other Vpx and Vpr orthologues shown to have transactivation activity in Fig 1e and Supplementary Fig 1f (HIV-2_ROD_ Vpx, SIV_MNE_027 Vpx, SIV_DRL_D3 Vpx, SIV_AGM_TAN1 Vpr, SIV_MND1_GB1 Vpr, and SIV_LST_524 Vpr) all decreased the levels of FAM208A (Supplementary Figs. 2b and c).

To assess whether disruption of FAM208A protein levels by Vpx was dependent upon the DCAF1 adaptor for the CUL4A/B ubiquitin ligase complex, as is the case for SAMHD1 (Sharova et al., 2008; Srivastava et al., 2008), Jurkat T cells were transduced with a lentivector that knocks down DCAF1 (Pertel et al., 2011a), or with a control knockdown vector. After selection with puromycin the cells were exposed for 18 hrs to SIV VLPs bearing Vpx, control VLPs that lacked Vpx, or no VLPs. In the DCAF1 knockdown cells, FAM208A protein levels were unchanged by Vpx, indicating that FAM208A disruption by Vpx was dependent upon DCAF1 (Fig. 2g).

Degradation of SAMHD1 requires direct interaction with Vpx or Vpr (Lim et al., 2012). To determine if Vpx similarly associates with proteins of the HUSH complex, HA-tagged FAM208A was co-transfected into HEK293 cells with FLAG-tagged SIV_MAC_251 Vpx or SIV_RCM_02CM8081 Vpx. When anti-FLAG antibody was used to immunoprecipitate either of the two Vpx proteins from the soluble cell lysate, HA-FAM208A was detected in the immunoprecipitate (Fig. 2h). The strength of the FAM208A signal in the Vpx pull-out increased when the co-transfected HEK293 cells were incubated with the proteasome inhibitor PR171, or when wild-type SIV_MAC_251 Vpx was replaced in the transfection by a mutant (Q76A) that is incapable of binding DCAF1 (Pertel et al., 2011a; Srivastava et al., 2008) (Fig. 2h and Supplementary Figs. 2d,e). These results demonstrate that FAM208A associates with Vpx and that the interaction results in proteasome-mediated degradation of FAM208A.

The experiments described above examined the effect of Vpx or Vpr on HIV-1 proviruses in which the reporter gene was transcribed by a heterologous promoter, either human EF1α, HSV TK, or the SFFV LTR (Figs 1 and 2, and Supplementary Fig. 1). To determine if Vpx is capable of activating a reporter gene driven by the HIV-1 LTR, the TNFα-responsive, J-Lat A1 clonal cell line was used (Jordan et al., 2003). In this experimental model of provirus latency, the HIV-1 LTR drives expression of a bicistronic mRNA encoding *tat* and *gfp* (Fig. 3a). Transduction with a lentivector expressing SIV_mac_251 Vpx, or knockdown of FAM208A, caused comparable increase in the percent GFP^+^ J-Lat A1 cells, whether the cells were stimulated with TNFα or not (Fig. 3b and c). Transduction of the J-Lat A1 cell line with lentivectors expressing *vpx* encoded by SIV_RCM_02CM8081 or SIV_MND2_5440, as well as with *vpr* encoded by SIV_MND1_GB1 or SIV_AGM_TAN1, caused similar increase in expression of the LTR-driven reporter gene (Supplementary Fig 3a).

**Figure 3.**
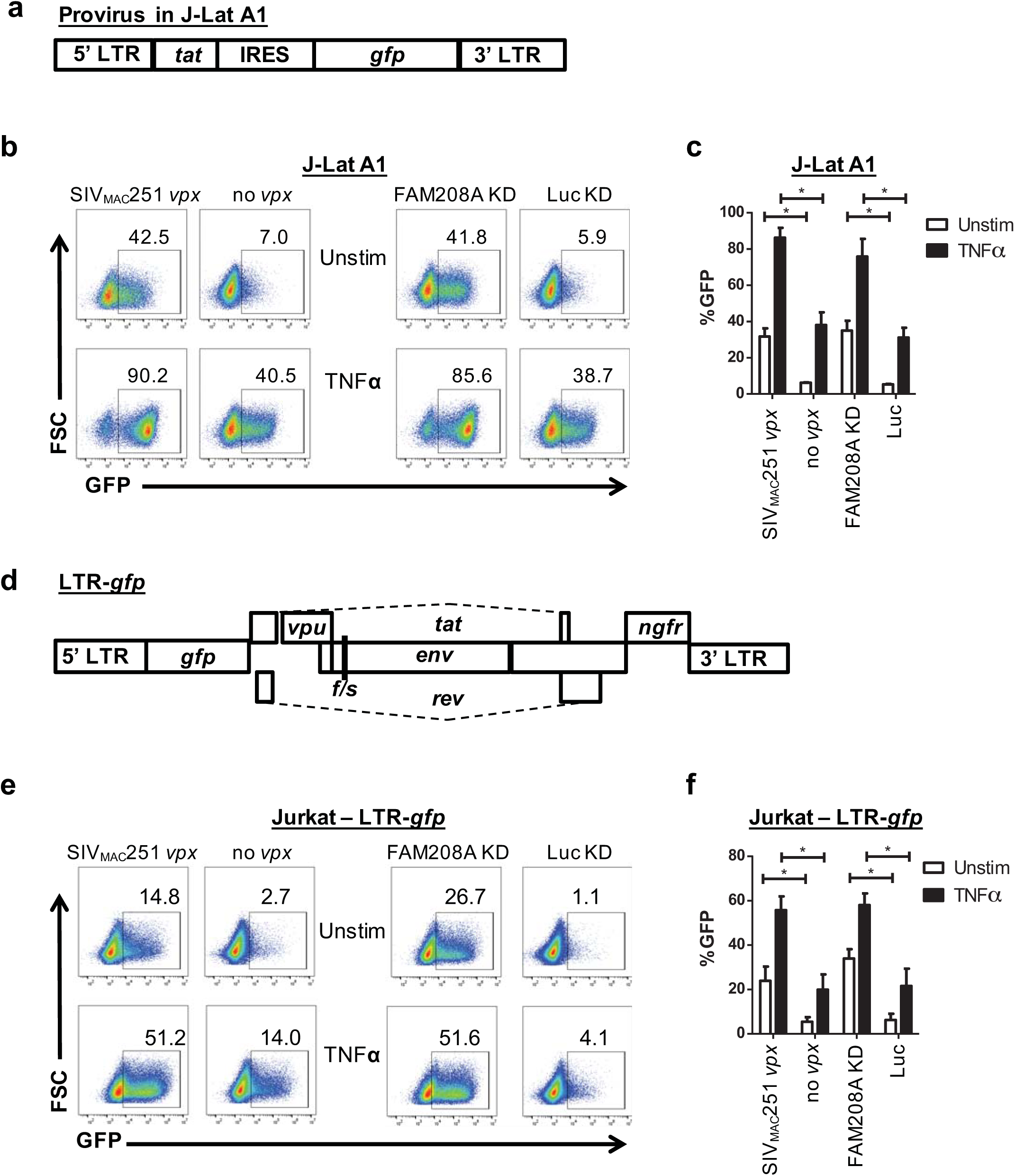
The HIV-1 LTR is activated by Vpx or disruption of FAM208A. **a,** Schematic of the HIV-1 minigenome integrated in the J-Lat A1 line. **b,** J-Lat A1 cells were transduced with Lenti 1 encoding SIV_MAC_251 *vpx* or Δ*vpx* control, or with lentivectors expressing shRNA targeting FAM208A or luciferase control. Transduced cells were selected with puromycin, and activated for 24 hrs with 10 ng/ml of TNFα. Representative GFP signal by flow is shown. **c,** Quantification of results from (**b**) and additional replicates (mean ± S.E.M., n=3 independent experiments). *, *P<0.02* **d,** Schematic of the LTR-*gfp* provirus used to analyze HIV-1 LTR driven *gfp* expression in pools of cells. **e**, Jurkat cells transduced with LTR-*gfp* were kept in culture for 4 wks and then transduced and assessed by flow cytometry, as in (**b**). **f,** Quantification of results from (**e**) (mean ± S.E.M., n=4 independent experiments) *, *P*<0.02

J-Lat A1 was selected to have a silent HIV-1 LTR-driven provirus with the ability to reactivate in response to TNFα (Jordan et al., 2003). The unique provirus within a clone such as J-Lat A1 may be sensitive to position-dependent silencing effects (Chen et al., 2016) and therefore may not accurately reflect the sensitivity of a population of HIV-1 proviruses to transcriptional activation by Vpx or to silencing by FAM208A. To address the effect of Vpx and FAM208A on on a population of proviruses with diverse integration sites, Jurkat T cells were transduced with an HIV-1 LTR driven reporter vector (LTR-*gfp)* that retains complete LTRs, *tat*, and *rev*, but has a frameshift mutation in *env*, an *ngfr* reporter gene in place of *nef*, and *gfp* in place of *gag, pol, vif, and vpr* (Fig. 3d). Four wks after transduction with LTR-GFP, the presence of latent proviruses within the pool of Jurkat cells was confirmed by reactivation with either TNFα or TCR-stimulation (Supplementary Fig. 3b). The Jurkat LTR-*gfp* cells were then transduced with vectors expressing SIV_MAC_251 Vpx or shRNA targeting FAM208A, and selected with puromycin. Compared with control cells, *vpx* or FAM208A knockdown increased the percentage of GFP^+^ cells, whether cells were treated with TNFα or not (Figs 3e and f). Similar results were obtained in three independently generated biological replicate experiments, in which *vpx* was delivered or FAM208A was knocked down, from four to eight wks after the first LTR-GFP transduction (Fig. 3f). Additionally, expression vectors for SIV_MND2_5440 Vpx, SIV_RCM_02CM8081 Vpx, SIV_MND1_GB1 Vpr, or SIV_AGM_TAN1 Vpr all increased GFP expression in Jurkat LTR-*gfp* cells (Supplementary Fig. 3c). Together, these experiments demonstrate that FAM208A contributes to the transcriptional repression of clonal or polyclonal LTR reporter lines, and that primate immunodeficiency viruses counteract this activity via their Vpx and Vpr proteins.

The effect of Vpx or FAM208A knockdown on spreading infection with replication-competent primate immunodeficiency viruses was tested next. Jurkat T cells transduced to express SIV_MAC_251 *vpx*, or cells transduced with control vector, were infected with HIV-1-ZsGreen, a replication-competent HIV-1_NL4-3_ clone, that encodes ZsGreen in place of *nef* (Supplementary Table 1). Infection was monitored by determining the percent ZsGreen^+^ cells with flow cytometry, every two days for ten days. Compared with the control, HIV-1 replication kinetics was accelerated by *vpx* (Fig. 4a). In similar fashion, HIV-1 infection of Jurkat cells transduced with the FAM208A knockdown vector resulted in faster replication kinetics (Fig. 4b).

**Figure 4.**
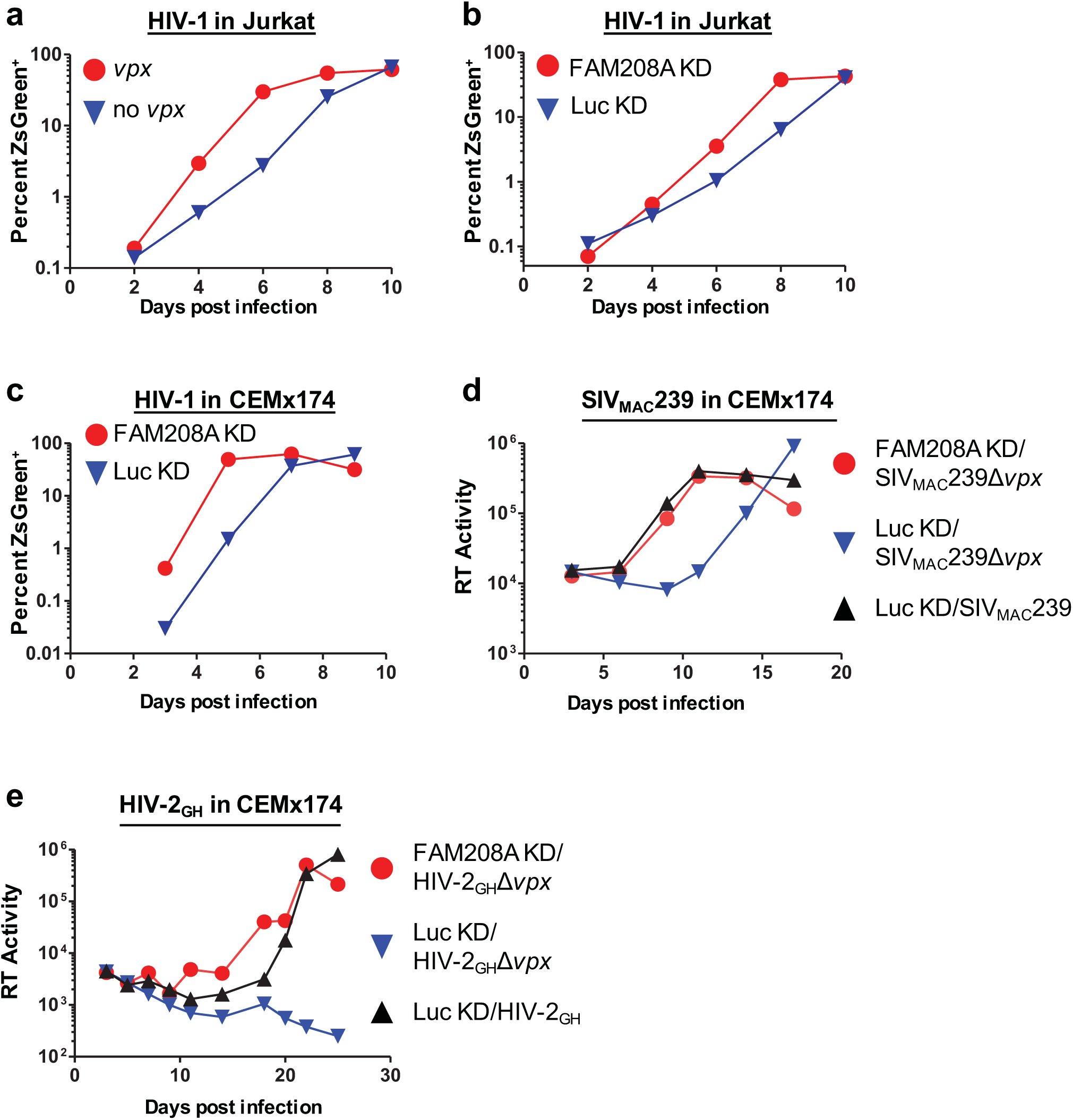
Vpx counteracts FAM208A restriction of HIV-1, SIV_MAC_239, or HIV-2_GH_, during spreading infection in CD4^+^ T cells. **a,b,** Replication of HIV-1-ZsGreen in Jurkat cells transduced with SIV_MAC_251 *vpx* or control (**a**), or with lentivectors expressing shRNA targeting FAM208A or Luc control (**b**). Replication kinetics was measured by flow cytometry for ZsGreen^+^ cells. **c,d,e**, Spreading infection of HIV-1-ZsGreen (**c**), SIV_MAC_239 or SIV_MAC_239Δvpx (**d**), and HIV-2_GH_ or HIV-2_GH_ ∆vpx virus in CEMx174 cells transduced with FAM208A or Luc control shRNA. Spread of HIV-1-ZsGreen was assessed by flow cytometry as in (**a**), while spread of SIV_mac_239 (**b**) and HIV-2_GH_ (**c**) was assessed by measuring the accumulation of reverse transcriptase (RT) activity in the supernatant. All data is representative of three repeat experiments.

HIV-1 vpr has no detectable effect on HIV-1 replication in tissue culture spreading infections with dividing target cells (Miller et al., 2017). This is presumably related to the cell cycle arrest toxicity (Chang et al., 2004; Goh et al., 1998; He et al., 1995; Re et al., 1995), and selection against *vpr* in tissue culture, since the effects of *vpr* on HIV-1 are evident when proviral expression is restricted to single cycle infection or cells are arrested with aphidicolin (Goh et al., 1998). Nonetheless, *vpr* offers a selective advantage in vivo since cloned *vpr* mutant virus was repaired when virus was injected into replication permissive chimps, or in an infected person (Goh et al., 1998).

SIV_MAC_239 does not replicate in Jurkat cells so CEMx174 cells were used to test the effect of FAM208A and *vpx* on replication of this virus. As in Jurkat cells, FAM208A knockdown increased HIV-1 replication kinetics in CEMx174 cells (Fig. 4c). Then, CEMx174 cells transduced with FAM208A or control knockdown vectors were challenged with SIV_MAC_239 or SIV_MAC_239-Δ*vpx* and replication was assessed by measuring reverse transcriptase activity in the supernatant. In the absence of *vpx*, SIV_MAC_239 replicated slower than the wild-type virus in control knockdown CEMx174 cells (Fig. 4d). This delay in SIV_MAC_239-Δ*vpx* replication kinetics was not observed when FAM208A was knocked down (Fig. 4d). Replication of HIV-2_GH_Δ*vpx* was undetectable in control knockdown CEMx174 cells (Fig. 4e). However, FAM208A knockdown rescued the replication of HIV-2_GH_Δ*vpx* to the level of wild-type HIV-2_GH_ in control cells (Fig. 4e). These experiments indicate that FAM208A inhibits primate immunodeficiency virus replication and that Vpx antagonizes this restriction, resulting in expression - or increased expression - from integrated proviruses, permitting virus spread.

The experiments reported here demonstrated that *vpx* and *vpr* activate transcription from silenced proviruses and that this activity was mimicked by knockdown of each of the HUSH complex components. These two observations were then shown to be linked by the finding that Vpx associated with, and promoted degradation of HUSH complex protein FAM208A, in a DCAF1- and proteasome-dependent manner. Latent provirus activation and human FAM208A degradation were exhibited by a broader range of primate immunodeficiency *vpx* and *vpr* orthologues than are capable of degrading human SAMHD1, perhaps due to the greater conservation and essential nature of FAM208A. Vpx and FAM208A disruption were important for transcriptional activation of latent HIV-1 provirus pools and for the ability of HIV-1, HIV-2, and SIV_MAC_ to effectively spread through cultured CD4^+^ T cells. Further understanding of the contributions of Vpx and Vpr and of the HUSH complex proteins, in concert with other transcriptional silencing mechanisms targeting HIV-1, is hoped to inform ongoing efforts to control or eliminate proviruses in HIV-1 infected patients.

## METHODS

### Plasmids

Sequences encoding 3xFLAG N-terminal-tagged Vpx and Vpr proteins were ordered as codon-optimized, gBlocks Gene Fragments (Integrated DNA Technologies; http://www.idtdna.com/) and cloned into either the pscALPS vector (Neagu et al., 2009) for transduction, or into pcDNA3.1 for transfection. pAPM-D4 is a truncated derivative of the pAPM lentivector (Pertel et al., 2011b) that expresses the puromycin acetyltransferase and miR30-based shRNA from the SFFV promoter. Supplementary Table 1 lists all plasmids used here, with corresponding addgene accession numbers, target sites used in particular knockdown vectors, and accession numbers for all the Vpx and Vpr orthologues tested here. All plasmid DNAs and sequences are available at https://www.addgene.org/Jeremy_Luban/.

### Cell culture

Cells were cultured at 37°C in 5% CO_2_ humidified incubators and monitored for mycoplasma contamination using the Mycoplasma Detection kit (Lonza LT07-318). HEK293 cells (ATCC) were used for viral production and were maintained in DMEM supplemented with 10% FBS, 20 mM L-glutamine (ThermoFisher), 25 mM HEPES pH 7.2 (SigmaAldrich), 1 mM sodium pyruvate (ThermoFisher), and 1x MEM non-essential amino acids (ThermoFisher). Jurkat and CEMx174 cells (ATCC) were cultured in RPMI-1640 supplemented with 10% heat inactivated FBS, 20 mM L-glutamine, 25 mM HEPES pH 7.2, 1 mM sodium pyruvate, 1x MEM non-essential amino acids and Pen/Strep (ThermoFischer) (RPMI-FBS complete). J-Lat A1 cells (Jordan et al., 2003) (NIH AIDS Reagent Program, catalogue #9852, donated by Eric Verdin) were cultured in RPMI-FBS complete media.

Leukopaks were obtained from anonymous, healthy, blood bank donors (New York Biologics, Southhampton, NY). As per NIH guidelines (http://grants.nih.gov/grants/policy/hs/faqs_aps_definitions.htm), experiments with these cells were declared non-human subjects research by the University of Massachusetts Medical School Institutional Review Board. PBMCs were isolated from leukopaks by gradient centrifugation on Histopaque-1077 (Sigma-Aldrich). CD4^+^ T cells were enriched from PBMCs using anti-CD4 microbeads (Miltenyi) and were >95% CD4^+^. CD4^+^ T cells were cultured in RPMI-FBS complete media in the presence of 50 U/mL hIL-2 (NIH AIDS Reagent Program, catalogue #136).

### Vector production

HEK293 cells were seeded at 75% confluency in 6-well plates and transfected with 6.25 μL Transit LT1 lipid reagent (Mirus) in 250 μL Opti-MEM (Gibco) with 2.25 µg total plasmid DNA. Full replicating virus was produced by transfection of 2.25μg of the indicated plasmid. Lenti-GFP reporters, LTR-GFP reporter, and shRNA lentivectors were produced by transfection of the lentivector, psPAX2 *gagpol* expression plasmid, and the pMD2.G VSV G expression plasmid, at a DNA ratio of 4:3:1. Vpx containing SIV-VLPs were produced by transfection at a 7:1 plasmid ratio of SIV3+ to pMD2.G, and ∆Vpx SIV VLPs were produced the same way using SIV3+ ∆Vpx plasmid. 12 hrs after transfection, media was changed to the specific media for the cells that were to be transduced. Viral supernatant was harvested 2 days later, filtered through a 0.45 µm filter, and stored at 4°C.

### Reverse Transcriptase assay

Virions in the transfection supernatant were quantified by a PCR-based assay for reverse transcriptase activity (Pertel et al., 2011b). 5 μl transfection supernatant were lysed in 5 μL 0.25% Triton X-100, 50 mM KCl, 100 mM Tris-HCl pH 7.4, and 0.4 U/μl RNase inhibitor (RiboLock, ThermoFisher). Viral lysate was then diluted 1:100 in a buffer of 5 mM (NH_4_)_2_SO_4_, 20 mM KCl, and 20 mM Tris–HCl pH 8.3. 10 μL was then added to a single-step, RT PCR assay with 35 nM MS2 RNA (IDT) as template, 500 nM of each primer (5’-TCCTGCTCAACTTCCTGTCGAG-3’ and 5’-CACAGGTCAAACCTCCTAGGAATG-3’), and hot-start Taq (Promega) in a buffer of 20 mM Tris-Cl pH 8.3, 5 mM (NH_4_)_2_SO_4_, 20 mM KCl, 5 mM MgCl_2_, 0.1 mg/ml BSA, 1/20,000 SYBR Green I (Invitrogen), and 200 μM dNTPs. The RT-PCR reaction was carried out in a Biorad CFX96 cycler with the following parameters: 42°C 20 min, 95°C 2 min, and 40 cycles [95°C for 5 s, 60°C 5 s, 72°C for 15 s and acquisition at 80°C for 5 s]. 3 part vector transfections typically yielded 10^6^ RT units/µL.

### Transductions

For generating pools of shRNA knockdown Jurkat and CEMx174 lines, cells were plated at 10^6^ cells/mL in RPMI-FBS complete and transduced with 10^7^ RT units of viral vector per 10^6^ cells, followed by selection with 1 μg/ml puromycin (InvivoGen, cat# ant-pr-1). To generate stable *gag-gfp* expressing Jurkat cells, cells were transduced as for shRNA KD above, followed by selection with 5 μg/mL blasticidin (InvivoGen, cat# ant-bl-1) at day 3 after transduction.

CD4^+^ T cells were stimulated in RPMI-FBS complete, with 50 U/ml IL-2 and 5 μg/mL PHA-P (Sigma, cat# L-1668). After 3 days, T cells were washed and replated at 3 × 10^6^ cells/mL in RPMI-FBS complete, with 50 U/ml IL-2. Cells were transduced with 10^8^ RT units of viral vector per 10^6^ cells followed by selection in 2 μg/mL puromycin.. After selection, cells were re-plated in RPMI-FBS complete with 50 U/ml IL-2 at 3 × 10^6^ cells/mL in RPMI-FBS complete and transduced again with the indicated GFP vectors, 10^8^ RT units of viral vector per 10^6^ cells. Transduced T cells were analyzed 4-5 days after the 2nd transduction.

### Lentiviral Infections

5 × 10^5^ Jurkat or CEMx174 cells were incubated with 5 × 10^7^ RT units of HIV-1_NL4.3_, HIV-2_GH_, HIV-2_GH_Δ*vpx*, SIV_MAC_239, or SIV_MAC_239Δ*vpx* virus stocks produced in HEK-293 cells for 12 hrs in RPMI-FBS complete media, followed by a wash in media and replated in 1 mL of media. Cells were split every 2-3 days and analyzed. For monitoring of HIV-1 ZsGreen infection, when cells were split, aliquots were fixed in BD Cytofix followed by analysis of GFP^+^ cells by flow cytometry to determine infection levels. For monitoring of SIV and HIV-2 infections, 50 μL aliquots of supernatant were analyzed for RT activity using the above described RT assay.

### Re-activation assays

LTR-driven GFP re-activation assays were performed with 10 ng/ml hTNFα (Invivogen, cat# rcyc-htnf), or with 1 μg/ml soluble α-CD3 and α-CD28 antibody. α-CD3 antibody (clone OKT3) and α-CD28 antibody (clone CD28.2) were provided by Lisa Cavacini (MassBiologics, Mattapan, Massachusetts).

### qRT-PCR

Total RNA was isolated from Jurkat cells using Trizol reagent followed by purification of RNA with RNeasy Plus Mini (Qiagen) with Turbo DNase (ThermoFisher) in order to limit DNA contamination. First-strand synthesis used Superscript III Vilo Master mix (Invitrogen) with random hexamers. qPCR was performed in 20 μL using SYBR green reagent (Applied Biosystems) with primers designed against *gag, gfp*, and *gapdh* for normalization.. Amplification was on a CFX96 Real Time Thermal Cycler (Bio-Rad) using the following program: 95°C for 10 min, then 45 cycles of 95°C for 15 s and 60°C for 60 s. Cells not transduced with Lenti-GFP vector were used as negative control and the housekeeping gene GAPDH was used to normalize expression levels. The primer sequences used were: *gag* primers (Forward: 5’- GCTGGAAATGTGGAAAGGAA-3’; Reverse: 5’-AGTCTCTTCGCCAAACCTGA-3’), *gfp* primers (Forward: 5’-GCAGAGGTGAAGTTCGAAGG-3’; Reverse: 5’- CCAATTGGTGTGTTCTGCTG-3’), *gapdh* primers (Forward: 5’- AGGGCTGCTTTTAACTCTGGT-3’; Reverse: 5’-CCCCACTTGATTTTGGAGGGA-3’).

### Flow cytometry

Cells were fixed in BD Cytofix Buffer prior to data acquisition on a BD C6 Accuri. Data was analyzed in FlowJo.

### Western Blot

Cells were washed in PBS, counted, normalized for cell number, and lysed directly in 1x SDS-PAGE sample buffer. Samples were run on NuPage 4-12% Bis-Tris gels followed by blotting onto nitrocellulose membranes. Primary antibodies used: FAM208A (Atlas, HPA00875), MPHOSPH8 (Proteintech, 16796-1-AP), PPHLN1 (Sigma, HPA038902), SETDB1 (Proteintech 11231-1-AP), DCAF1 (Proteintech, 11612-1-AP), FLAG (Novus, NB600-345), FLAG (Sigma, F1804, used for IP), and HA (Biolegend, 901501).

### Vpr and Vpx phylogeny

The following Vpr and Vpx amino acid sequence alignments were obtained from the Los Alamos National Laboratories (LANL) HIV sequence database: 2016 HIV-1/SIVcpz Vpr, 2016 HIV-2/SIVsmm Vpr, 2016 HIV-2/SIVsmm Vpx, 2016 other SIV Vpr, and 2016 other Vpx. Consensus sequences were generated for HIV-1 group M subtypes A, B, C, D, F, G, H, I, J, and those designated U in the LANL database, as well as group N. A master alignment was scaffolded from the above alignments and re-aligned by hand. Redundant SIV and HIV-2 Vpr and Vpx sequences were removed, and the sequences of individual HIV-1 isolates were replaced with the consensus sequences. This was used to generate a master phylogeny using RAxML 8.2.11, as implemented in Geneious with gamma LG substitution model and Rapid Bootstrapping with search for best scoring tree algorithm. This master tree was utilized to identify major relationships and identify a reduced number of sequences to retain while maintaining the overall phylogenic structure. Vpx and Vpr sequences from the following viral isolates were retained: HQ179987, L20571, M15390, AF208027, AB731738, KP890355, M15390, AF208027, AB731738, KP890355, U58991, M30931, L40990, KJ461715, AF301156, U42720, AY169968, DQ373065, DQ373064, DQ374658, FJ919724, AJ580407, KM378563, KM378563, FJ424871, M66437, AF468659, AF468658, AF188116, M76764, LC114462, M27470, AY159322, AY159322, U79412, U79412, AY340701, AY340700, EF070329, KF304707, FM165200, HM803690, HM803689, AF382829, AF349680, HM803690, HM803689, AF349680, U04005, JX860432, JX860430, JX860426, JX860432, M83293, M83293, AF131870, AY523867, AM182197, AM713177, U26942, and the HIV-1 group M clade B consensus. These sequences were used to generate a phylogeny using the same method as above. Superfluous taxa were pruned from this phylogeny using Mesquite 3.4 and the resulting tree was visualized in FigTree v1.4.3.

### Sampling

At least three biological replicates were performed for all experiments. The screen for factors mediating silencing of the Lenti-GFP vector utilized 3 target sequences for each candidate gene. Flow cytometry plots in the figures show representative data taken from experiments performed at the same time. HIV-1, HIV-2, and SIV spreading experiments were repeated 3 times each and representative data of one such experiment is shown.

### Statistics

Information regarding the statistical tests utilized, and the n values, are found in the figure legends. Statistical analysis of the knockdown screen of factors involved in silencing of Lenti-GFP was analyzed by one-way ANOVA with Dunnett post test comparing 3 shRNA target sites to control knockdown conditions. All statistics presented were performed using PRISM 5.0 (GraphPAD Software, La Jolla, CA).

## ACKNOWLEDGEMENTS

The authors wish to dedicate these experiments to the memory of Jan Svoboda (1934-2017), whose demonstration that cells may carry Rous sarcoma virus genetic information in the absence of any infectious virus production provided support to the proviral hypothesis. We thank Lisa Cavacini for anti-CD3 and anti-CD28 antibodies, and Akio Adachi and Mikako Fujita for pGL-St and pGL-An. The following reagents were obtained through the AIDS Reagent Program, Division of AIDS, NIAID, NIH: J-Lat Tat-GFP Cells (A1) from Dr. Eric Verdin, and SIV_mac_239 SpX and SIV_mac_239 SpX ΔVpx from Dr. Ronald C. Desrosiers.

## Author Contributions

L.Y. and J.L. designed the experiments. L.Y. performed the experiments with assistance from M.H.G., K.K., S.L.G., S.M., A.D., and W.E.D., L.Y. and J.L. analyzed the experimental data. All authors contributed to the writing of the manuscript.

## Data and materials availability

all data needed to evaluate the conclusions in the paper are present in the paper or in the supplementary table. The plasmids described in Supplementary Table 1, along with their complete nucleotide sequences, are available at https://www.addgene.org/Jeremy_Luban/. Correspondence and requests for materials should be addressed to J.L.

